# DNA Preparation in *Vitis vinifera* L. For Third Generation Sequencing

**DOI:** 10.1101/2021.03.11.435003

**Authors:** Bibi C. Androniki, Kollias Anastasios, Astrinaki Maria, Vassou Despoina, Kafetzopoulos Dimitris, Kalantidis Kriton, Moschou N. Panagiotis

**Author notes:** Coresponding author: Bibi C. Androniki. **Contact for reagents and resource sharing**, Further information and requests for resources and reagents should be directed to and will be fulfilled by the Coresponding author, Bibi C. Androniki, Address: Institute of Molecular Biology & Biotechnology, Foundation for Research & Technology - Hellas, Nikolaou Plastira 100 GR-70013, Heraklion, Crete GREECE. **Author Approvals:** All authors have seen and approved the manuscript, and that it hasn’t been accepted or published elsewhere.

## Abstract

**Background:** There have been several attempts to sequence the genome of *Vitis vinifera* L. (grapevine), utilizing low-resolution second-generation platforms. Nevertheless, the characterization of the grapevine genetic resources and its adaptation to vulnerable conditions could be better addressed through extensive and high-resolution genome sequencing.

MinION is a third-generation sequencer preferred by many laboratories due to its relatively low cost, ease of use and small size. Even though this long-read technology has been rapidly improving, to reach its full potential requires high-quality DNA.

**Results:** Here we establish a workflow for DNA extraction suitable for MinION sequencing long reads from grapevine. This protocol was tested with leaf samples from different positions on annual growing branches of grapevine, Purified nuclei from fresh young leaves that led to high quality, long DNA fragments, suitable for long-read sequencing were successfully generated. It is evident that longer reads in grapevine associate with both fresh tissue and adjusted conditions used for nuclei purification.

**Conclusions:** We propose that this workflow presents a significant advancement for long-read quality DNA isolation for grapevine and likely other plant species.

## Background

Grapevine (*Vitis vinifera* L.) is one of the worlds’ most important commodity crops, mainly used for the production of wine, table grapes and dried grapes, although other minor applications are also widespread in some regions. Grapevine is a model organism for the study of perenniality and non-climacteric fruit ripening (Cantu and Walker, 2019). The first grapevine genome was released by the French-Italian Public Consortium for Grapevine Genome Characterization in 2007 (Jaillon *et al.*, 2007). The sequence for this first version was obtained using a whole-genome shotgun strategy and the Sanger sequencing technology, while later was refined through a 4X additional coverage (Adam-Blondon, 2014). Finally, grapevine reference genome assembly (12X.v2) was further refined by Illumina HiSeq 2500 sequencer (Illumina Inc.). Despite the technical advances, the short-reads produced by the second-generation platforms cannot resolve complex genomic regions which limit the characterization of genetic resources and detection of genomic structural variants (SVs) (Alkan et al., 2011, Pollard et al., 2018)

Third-generation sequencing platforms achieve an unprecedented combination of speed, accuracy, and flexibility. These platforms include single-molecule real-time sequencing (SMRT) and nanopore techniques enabling highly-parallelized, amplification-free sequencing of individual DNA molecules, thereby enabling long reads up to hundreds of kilobases (kb). This feature is useful for resolving highly repetitive genome features make it possible to generate phased de novo genome assemblies with greater continuity and completeness. MinION is a 3^rd^ gen sequencer from Oxford Nanopore Technologies (ONT) that enables real-time analysis of sequence data on a regular computer. The MinION uses nanopores to sequence a single DNA molecule per pore; this has significant potential advantages over the current widely used technologies (Ion Torrent, Illumina), which rely on sequencing clusters of amplified DNA molecules.

A standard MinION flow cell can yield up to 40 Gb of data within 72 h and enough sequencing depth can be achieved within a few minutes (Butt et al., 2018). ONT has produced a flow cell called “Flongle” (R9.4.1 nanopores), which is smaller and cheaper than regular flow cell (SpotON flow cell Mk I, R9.4.1 nanopores) (Grädel *et al.*, 2019). Flongle is mounted onto a Flongle adapter that contains the ONT proprietary sensor array and is compatible with MinION. A Flongle harbours 126 sequencing channels instead of the standard 512, is typically designed for a single sample instead of multiple ones and is thus used to rapidly assess sample quality before scaling up.

Here we establish a workflow for DNA extraction compatible with MinION sequencing for long reads from grapevine leaves. To fully benefit from nanopore sequencing platforms by obtaining longer sequencing read lengths, DNA fragmentation needs to be minimized. Currently, sample preparation still requires optimization before sequencing, which can be challenging. In grapevine, the yield and quality of DNA can be significantly compromised by polyphenols, polysaccharides, and proteins, which are abundant during different stages of berry and leaf development (Iandolino *et al.*, 2004). ONT proposed a method to obtain long-read sequencing by integrating nuclei isolation with Nanobind DNA isolation (Circulomics Inc.) (Workman *et al.*, 2018). Nanobind is a magnetic disk covered with a high-density micro and nanostructured silica. This technology yields high molecular weight genomic DNA (50–300 kb) from nuclei; it was successfully used in *Sequoia sempervirens* and *Zea mays* (Workman *et al.*, 2018). Our main objective here was to optimize the workflow for grapevine, to get high quality, long DNA fragments, suitable for long-read sequencing. We foresee that the pipelines we suggest herein will apply to a large number of species.

## Methods

### Plant Material

Leaf samples from different positions on annually growing branches of *Vitis vinifera* L. plants were collected from field-grown plants, at the IMBB campus at 10 am. Some samples were store at −80°C, while others were used directly for DNA extraction.

### DNA extraction

DNA was extracted as described by Workman et al. (2018) with minor modifications using fresh leaves from the upper nodal positions of annually growing branches. One gram of fresh leaf-tissue from the uppermost nodal position was used for intact nuclei isolation. Leaf tissues were ground in liquid N2 with a mortar and pestle, for at least 30 min. The ground tissue is transferred to a tube where 10 mL of fresh nuclear isolation buffer (NIB: prepared according to Workman et al. 2018) was added. Nuclei were isolated in 10 ml NIB. Tubes were agitated on a rotary platform at max speed for 15 min at 4 °C. Homogenates were filtrated through a strainer (40nm) and then through 5 layers of Miracloth into 50 mL tubes. Samples were centrifuged at 4 °C for 20 min at varying centrifugal forces (herein 2500g, 4000g, and 5000g). The supernatants were discarded and 1 mL cold NIB was added to each pellet, and then resuspended with a paintbrush presoaked in NIB; suspensions were transferred in a new tube. Volumes were adjusted to 15ml by ice-cold NIB and the samples were centrifuged at 5,000g, 2,500g or 4,000g for 15 min at 4 °C. This step was repeated 3-4 times until the supernatants were colourless. Next, pellets were resuspended in 1 mL 1X Homogenization Buffer (HB: prepared according to Workman et al. 2018) and transferred to 1.5ml microcentrifuge tubes. Samples were either snap frozen or used directly. For long term storage, samples were spun down at 2,500 g for 5 min, supernatants were discarded and pellets were snap-frozen in liquid N2 and stored at −80 °C. Nuclei were stained for 5 min with an aqueous solution of 1 μm DAPI (4’,6-Diamidine-2’-phenylindole dihydrochloride) and examined by Confocal Microscopy.

### Confocal Microscopy

For confocal microscopy of nuclei, fluorescence images were captured on a Leica SP8 (Leica Camera AG, Wetzlar, Germany). The fluorescent images (micrographs) were acquired at RT with 40x objective (N.A.=1.2), pinhole adjusted to 1 airy unit, and mounting medium ddH_2_O. The excitation wavelength was 405nm (Emission 412nm-467nm) for DAPI.

### Nanobind-assisted DNA purification

Pellets were resuspended in 60 μL of Proteinase ? and vortexed, followed by addition of 20 μL RNase A and vortexed again until a homogenous mixture was obtained. Then, 140 μL Buffer PL1 was added and vortexed extensively. Thorough mixing at this step is necessary to ensure complete lysis of the nuclei. Samples appeared cloudy and very viscous but homogeneous and were incubated on an Eppendorf ThermoMixer^TM^ C (Thermo Fisher Scientific Inc.), at 55 °C and 900 rpm for 30 min. Lysates were centrifuged for 5 min at 16,000*g* at RT. The supernatants were transferred to 1.5 mL Protein LoBind microcentrifuge tubes using a wide bore pipette. Nanobind disks were added to the supernatants followed by addition of a 1X volume of isopropanol. The samples were carefully mixed by manual inversion for 20 min at RT. The microtubes were placed in a magnetic tube rack as per instructions from the Nanobind Plant Nuclei Big DNA Kit. Supernatants were discarded, 500 μL of Buffer PW1 were added and samples were mixed 4X by inversion. This step was repeated twice then, any residual liquid from the tube caps was removed by spinning on a microcentrifuge for 2s. Finally 100-200 μL Buffer EB was added and the tubes were spun on a microcentrifuge for 2 s and incubated at RT for 10 min. The extracted DNA was collected by transferring the eluate to a 1.5 mL microcentrifuge tube using a wide bore pipette. Tubes were further spun on a microcentrifuge for 5 s and remaining eluted DNA was collected and transferred to the same 1.5 mL microcentrifuge tube. The quantity and the quality of the extracted DNA were assessed by NanoDrop (NanoDrop ND-1000 Spectrophotometer) and Qubit (Invitrogen Qubit 3’).

### DNA Library construction For Flongle

According to the manufacturer, 3 to 20 fmoles of DNA are required for Flonge Cells. Libraries were prepared using the 1D ligation sequencing kit (SQK-LSK109). Briefly, the workflow consisted of first an end-repair/dA-tailing step to repair blunt ends and to add an “Adenine” to the 3 ‘ end of the amplicon, followed by adapter and motor protein ligation onto the prepared ends using NEB ligase (New England Biolabs). Subsequently, a bead-based purification step was used to enrich the adapter-ligated fragments and remove excessive nucleotides and enzymes. Finally, the library mixture was loaded into the Flongle flow cell and the MinKNOW GUI (version 19.06.8) was used for data acquisition as per recommendation from ONT. Each sequencing experiment was run for 24 hours (>60 active pores) or after a minimum of 500 reads were acquired for downstream data analysis. Promega Biomath Calculator (https://worldwide.promega.com/resources/tools/biomath/) was used to determine the amount of DNA in fmoles, using the starting yield of nucleic acids and the estimation of N50 of the reads ca. at 30-40 kb. The final yield of the libraries varied between 3-5 fmoles, which is the minimum amount required. Therefore, for the construction of the second library (referred to as “Vitis 2”, 4,000g), the starting yield of DNA was increased 4x at 2.5μg. After the ligation with the reagents used in double quantities, the final yield was 0.63μg, with 25% recovery, and was loaded in the Flongle.

### In Silico Data Analyses

Raw FAST5 files produced by MinION were base-called using the integrated live base-calling setting in the MinKNOW Core 3.6.5. and Guppy 3.2.10 (Wick, Judd and Holt, 2019) that converts Fast5 files to FastQ. To proceed with the analysis it is important to know that the quality baseline (Q) is 7 by default in MinION. The goal is to achieve Q ≥ 8.5 and increased mean read length and higher outcome. Control DNA was eliminated by NanoLyse [1.2.0] and served as quality control for each run using NanoPlot1.13.0, and in combination using NanoComp 0.16.4. These tools are integrated into the NanoPack suite (De Coster *et al.*, 2018).

## Results

### Workflow for DNA extraction

To establish an efficient protocol for grapevine DNA purification compatible with 3^rd^ generation sequencing, we tested different protocols for nuclei isolation. As initial attempts to purify nuclei at standard low (2,000g) or high (6,000g) relative centrifugal forces (*g*=timesxEarth’s gravitational force) did not yield nuclei, we assumed that the parameter *g* may be a limiting factor when it comes to nuclei purification and should be experimentally determined. Nuclei from different species, organs, tissues and cell ages show significant differences in their size and ploidy and thus mass. We thus further tried three different nuclei purification schemes, with variable centrifugal force (i.e. 2,500, 4,000 (“*Vitis 2*”) and 5,000 g (“*Vitis 1*”). We determined that the optimal scheme for nuclei purification was Vitis 2, while *Vitis 1* yielded nuclei that were mechanically perturbed (**Fig. 1**); at 2,500g we obtained very few nuclei. Furthermore, Vitis 2 ratios of 260/280 and 260/230 in NanoDrop were higher, compared to *Vitis 1,* showing that *Vitis 2* DNA was of better quality (**Table 1**). The DNA extraction took us on average 3-4 h for nuclei preparations and 2-3 h for Nanobind-assisted DNA purification. The workflow with the duration of each step is presented in more detail in **Fig. 2**.

**Fig 1.**
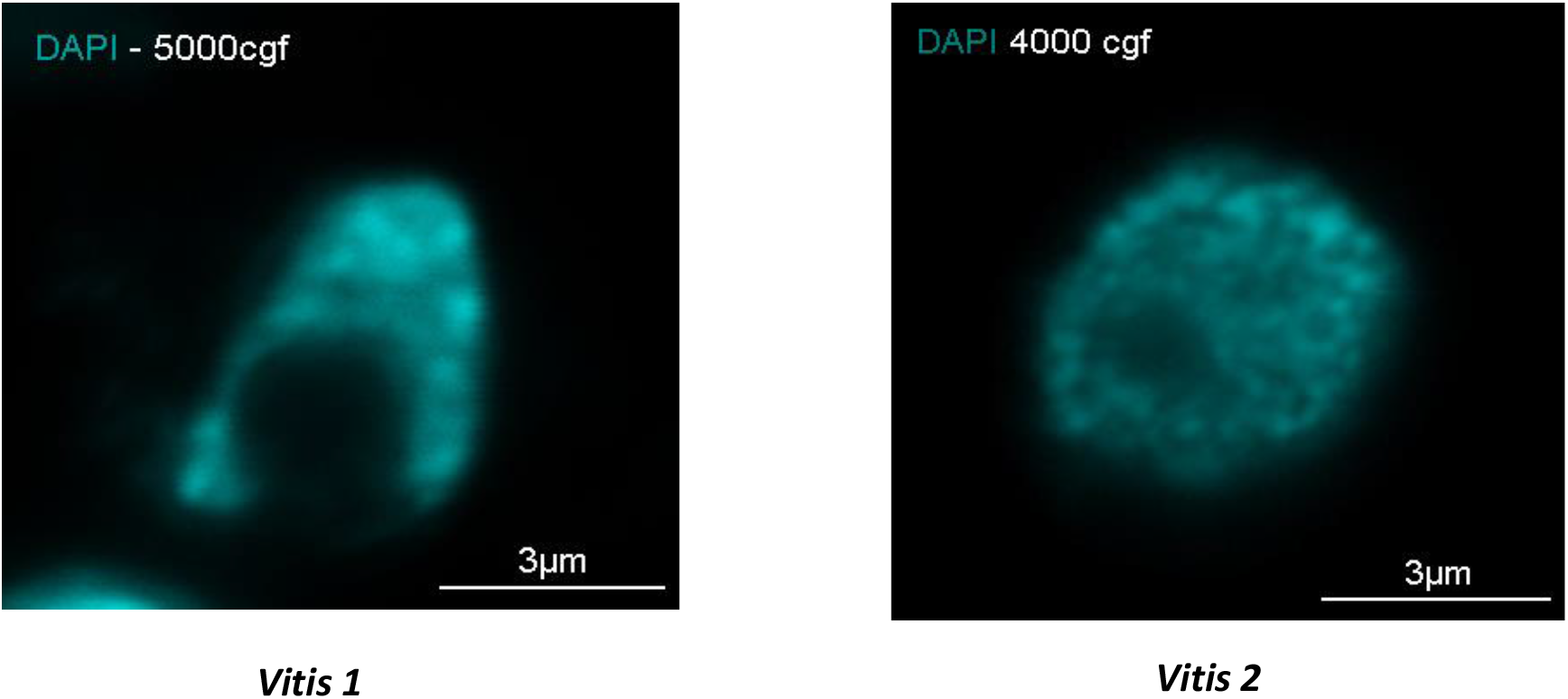
Confocal micrographs of nuclei isolated from grapevine leaves and stained by DAPI. Left: *Vitis 1* at RCF of 5,000g; Right: *Vitis 2* at RCF of 4,000g. Data in (A) and (B) are representative images of an experimental replicated three times. Scale bars are shown.

**Fig 2.**
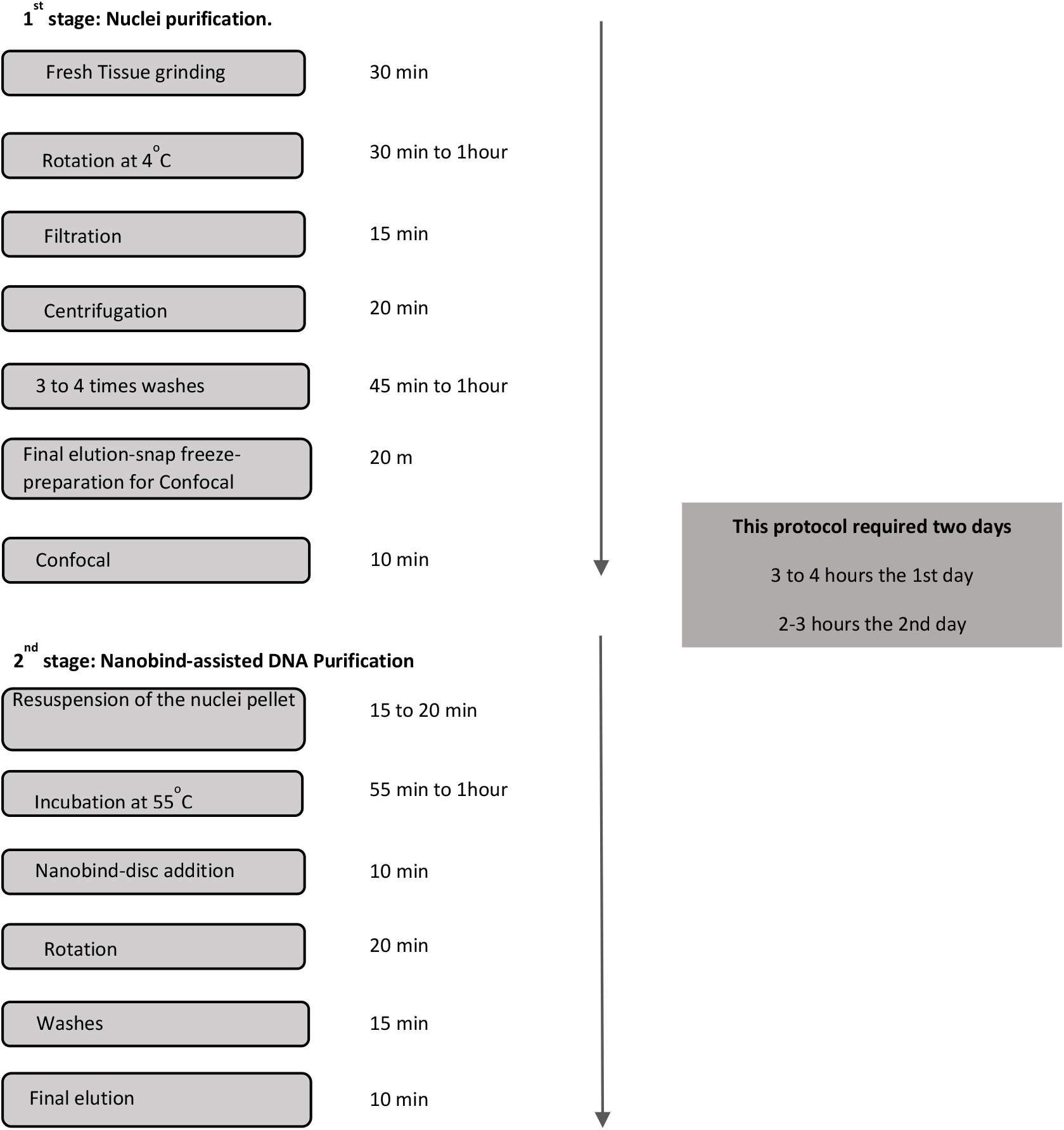
Work flow for DNA extraction

**Table 1.** DNA concentration (ng/μL) and the ratios 260/280, 260/230 from NanoDrop of Vitis 1 and Vitis 2 samples.

| ID of sample | Yield (ng/μl) | Ratio 260/280 | Ratio 260/230 | Qubit (ng/μl) |
| --- | --- | --- | --- | --- |
| *Vitis_1* | 472.7 | 1.7 | 1.43 | 375 |
| *Vitis_2* | 292 | 1.82 | 1.75 | 397.5 |

### ONT-Flongle flow cell sequencing results

The increased amount of DNA loaded in library preparation of *Vitis 2* and the difference in the extraction conditions were reflected by the output of the total bases that reached approximately 672 Mb in the *Vitis 2* run. Mean read length from 3,277.6 in *Vitis 1* increased to 5,464.8 in Vitis 2. Similarly, mean read quality increased in *Vitis 2* to 9.3 from 7.3 in *Vitis 1.* The distributions of read length and the read quality are shown in **Table 2** and **Fig.3**.

**Fig 3.**
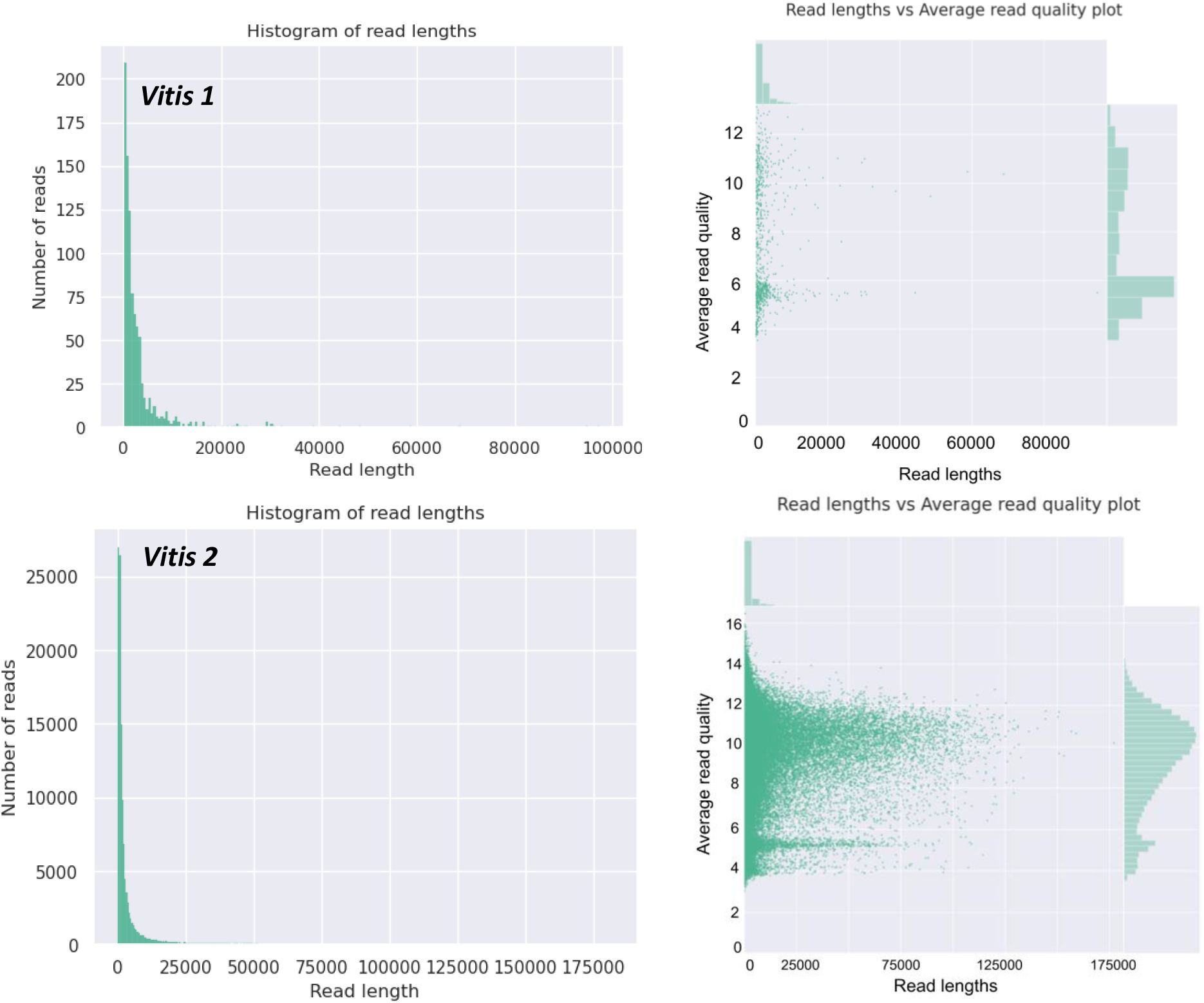
Distribution of reads obtained and the average read quality of *Vitis 1* and *Vitis 2.*

**Table 2:** NanoComp analysis of the two MinION sequencing runs. Statistics of Nanopore sequencing data without the cDNA sequencing

| **Nanopore Statistics** |  |  |
| --- | --- | --- |
|  | ***Vitis_1*** | ***Vitis_2*** |
| Reads number | 921.0 | 122,987.0 |
| Mean reads length | 3,277.6 | 5,464.8 |
| Reads length N50 | 7,463.0 | 29,948.0 |
| Mean read quality | 7.3 | 9.3 |
| Quality filtered reads (> Q7) | 404 (43.9%) 1.4Mb | 102458 (83.3%) 595.3Mb |
| Total bases | 3,018,681.0 | 672,098,346.0 |

Next, we evaluated the performance of both sequencing runs. To this end, we performed read counts, read length-average base-call quality score comparison, and total Gbs sequenced for comparing the two runs, *Vitis 1* and *Vitis 2* (**Fig.4**). From these results, it is evident that longer reads in grapevine associate with both fresh tissue and adjusted RCF.

**Fig 4.**
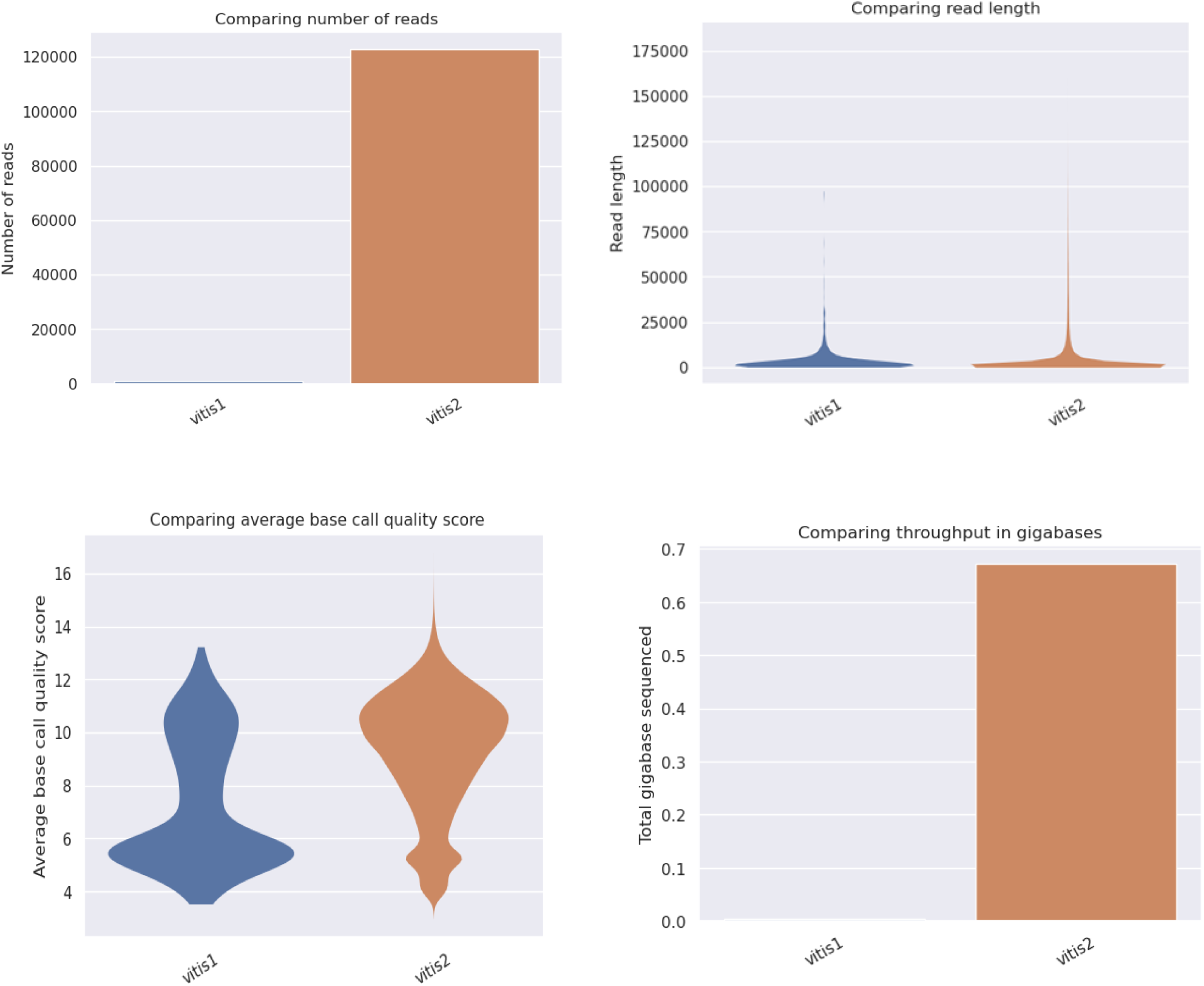
Comparison between reads number produced between *Vitis* 1 and 2. (A) Number of reads of *Vitis* 1 and 2. (B) Volcano plots of read lengths of *Vitis* 1 and 2. (C) Volcano plots showing average lengths and base call quality score between the two runs. (D) Total Gbs acquired between the two runs.

## Discussion

We have established a pipeline for DNA purification from grapevine, with the ability to yield high quality DNA for 3^rd^ generation sequencing. This pipeline efficiently incorporates circulomic, flonge and bioinformatics. Our results imply that the optimal centrifugal force used for the nuclei purification in circulomics should be experimentally determined and adjusted according to the stage, tissue type and age. Young leaf tissue was the optimal DNA source possibly because it has more cells per unit-area, and typically contains less secondary metabolites and phenolics (Murray & Thompson, 1980, Doyle & Doyle, 1987, Peterson et al., 1997, Iandolino et al., 2004). Likewise, we failed to obtain DNA from old leaves. Regarding storage conditions, we extracted DNA from freshly collected tissue and tissue immediately frozen in liquid N2 and stored at −80 °C to avoid degradation by nucleases (Ribeiro & Lovato, 2007, Varma et al., 2007; Knebelsberger & Stöger, 2012). However, we observed that even young leaves should be used only fresh, as otherwise, the DNA quality dropped dramatically. Samples stored for >3 months at −80 °C did not yield nuclei and thus DNA. Likewise, in human studies, read lengths were longer when the DNA was freshly extracted (Jain *et al.*, 2018).

We evaluated the performance of the yielded DNA using Flonge technology. Flongle is an adapter for MinION or that enables direct, real-time DNA sequencing single-use flow cells and is the quickest, most accessible solution for smaller tests and experiments (up to 2 Gb). From our results, it is evident that longer reads in grapevine, which can reach up to 50 kb, associate with both fresh tissue and centrifugal force. This study was an attempt to perform several modifications to a DNA extraction protocol described by Workman et al. (2018), to yield high-molecular-weight DNA for grapevine.

## Conclusion

In conclusion, Minion platform would require optimization of the plant material to achieve proper DNA quality. Our workflow could be adapted for the isolation of high quality, long-read-ready DNA from various plant species.

## Acknowledgements

This research has been co-financed by the European Union and Greek national funds through the Operational Program Competitiveness, Entrepreneurship and Innovation, under the call RESEARCH – CREATE – INNOVATE (project code:T1EDK—04363. CloseViva: A Collaborative Network for the Exploitation and Clonal Selection of Greek Vine Varieties and Valorization of the Genetic Material). In addition, this research has been financed by the project “Development of national research network to «Grape», Division 1. (Development of national research network to «Olive», to «Grape», to «Honey», and to «Livestock»)”, with the code 2018ΣE01300000 of the National Scale of the Public Investments Program of the General Secretariat of Research and Innovation.

## Authors’ contributions

Study conception and design: ACB, PNM, DK

Acquisition of data: ACB, AK

Analysis and interpretation of data: ACB, AM, DV

Drafting of manuscript: ACB, PNM, KK

Critical revision: ACB, PNM, KK

## References

Adam-Blondon, A.-F. (2014) ‘GRAPEVINE GENOME UPDATE AND BEYOND’, Acta Horticulturae, 1046(1046), pp. 311–318. doi: 10.17660/ActaHortic.2014.1046.42.

Alkan, C., Coe, B. P. and Eichler, E. E. (2011) ‘Genome structural variation discovery and genotyping’, Nature Reviews Genetics, 12(5), pp. 363–376. doi: 10.1038/nrg2958.

Cantu, D. and Walker, A. (Eds.. (2019) The Grape Genome. Edited by D. Cantu and M. A. Walker. Cham: Springer International Publishing (Compendium of Plant Genomes). doi: 10.1007/978-3-030-18601-2.

De Coster, W. et al. (2018) ‘NanoPack: Visualizing and processing long-read sequencing data’, Bioinformatics, 34(15), pp. 2666–2669. doi: 10.1093/bioinformatics/bty149.

Doyle, J. J. and Doyle, J. L. (1987) ‘Doyle_plantDNAextractCTAB_1987.pdf’, Phytochemical Bulletin, pp. 11–15. Available at: https://webpages.uncc.edu/~jweller2/pages/BINF8350f2011/BINF8350_Readings/Doyle_plantDNAextractCTAB_1987.pdf.

Grädel, C. et al. (2019) ‘Rapid and Cost-Efficient Enterovirus Genotyping from Clinical Samples Using Flongle Flow Cells’, Genes, 10(659), pp. 1–12.

Iandolino, a B. et al. (2004) ‘Libraries From Grapevine (Vitis vinifera L.)’, Plant Molecular Biology Reporter, 22(September), pp. 269–278.

Jaillon, O. et al. (2007) ‘The grapevine genome sequence suggests ancestral hexaploidization in major angiosperm phyla’, Nature, 449(7161), pp. 463–467. doi: 10.1038/nature06148.

Jain, M. et al. (2018) ‘Nanopore sequencing and assembly of a human genome with ultra-long reads’, Nature Biotechnology, 36(4), pp. 338–345. doi: 10.1038/nbt.4060.

Knebelsberger, T. and Stöger, I. (2012) ‘DNA extraction, preservation, and amplification’, Methods in Molecular Biology, 858, pp. 311–338. doi: 10.1007/978-1-61779-591-6_14.

Murray, M. G. and Thompson, W. F. (1980) ‘Rapid isolation of high molecular weight plant DNA’, Nucleic Acids Research, 8(19), pp. 4321–4326. doi: 10.1093/nar/8.19.4321.

Peterson, R. A., Balasubramanian, S. and Bronnenberg, B. J. (1997) ‘Exploring the implications of the internet for consumer marketing’, Journal of the Academy of Marketing Science, 25(4), pp. 329–346. doi: 10.1177/0092070397254005.

Pollard, M. O. et al. (2018) ‘Long reads: their purpose and place’, Human molecular genetics, 27(R2), pp. R234–R241. doi: 10.1093/hmg/ddy177.

Ribeiro, R. A. and Lovato, M. B. (2007) ‘Comparative analysis of different DNA extraction protocols in fresh and herbarium specimens of the genus Dalbergia’, Genetics and Molecular Research, 6(1), pp. 173–187.

Varma, A., Padh, H. and Shrivastava, N. (2007) ‘Plant genomic DNA isolation: An art or a science’, Biotechnology Journal, 2(3), pp. 386–392. doi: 10.1002/biot.200600195.

Wick, R. R., Judd, L. M. and Holt, K. E. (2019) ‘Performance of neural network basecalling tools for Oxford Nanopore sequencing’, bioRxiv, pp. 1–10. doi: 10.1101/543439.

Workman, R. et al. (2018) ‘PROTOCOL EXCHANGE | COMMUNITY CONTRIBUTED High Molecular Weight DNA Extraction from Recalcitrant Plant Species for Third Generation Sequencing’, pp. 1–12. Available at: https://www.nature.com/protocolexchange/protocols/6785.

